# Anodal tsDCS restores the structure and function of the disrupted proprioceptive Ia synapses on spinal motoneurons in the SOD1 G93A mouse model of ALS

**DOI:** 10.1101/2024.08.02.606278

**Authors:** T. Jankowiak, M. Cholewiński, K. Kryściak, E. Krok, K. Grycz, M. Bączyk

## Abstract

An imbalance between cells’ intrinsic excitability and synaptic excitation levels is the basis of spinal motoneuron (MN) pathophysiology in Amyotrophic Lateral Sclerosis. Recently, a restoration of the deficient Ia synaptic excitation of spinal MNs was achieved by applying acute trans-spinal direct current stimulation (tsDCS) to presymptomatic SOD1 G93A mice. Here we investigate whether two-week repeated tsDCS applied to presymptomatic SOD1 animals can provoke spinal MN neuroplasticity and reduce the disease burden. Anodal, cathodal or sham polarisation of 100 µA was applied to P30-P35 SOD1 G93A mice; passive membrane properties and Ia excitatory post-synaptic potential (EPSP) characteristics were investigated by intracellular recordings of spinal MNs in vivo. A second cohort of polarized animals was used to test the impact of our intervention on Ia synapse morphology, MN intracellular metabolic pathways activity, and disease markers. Anodal tsDCS evoked a strong increase in maximal Ia EPSPs, coupled with a significant upregulation of vesicular glutamate transporter levels and GlurR4 subunits of AMPA receptors at the Ia synapse. On the other hand, cathodal polarisation failed to induce any significant alteration to Ia synapse morphology but did increase both peak and plateau input resistance and recovered the abnormal paired-pulse ratio. Unexpectedly, the changes in MN electrophysiological profile and Ia synapse morphology did not translate into alterations of intracellular pathways ctivity and did not decrease the disease burden. Altogether our results indicate a strong polarity-dependent plasticity of spinal MNs in SOD1 G93A mice in response to tsDCS, which nevertheless appears insufficient to alter disease dynamics.

**Highlights:**

- 14-days of trans-spinal direct current stimulation (tsDCS) alters the electrophysiological properties and morphology of Ia proprioceptive synapses on spinal MNs in SOD1 G93A mouse model of ALS
- Anodal (depolarising) tsDCS increases MN synaptic excitation and restores the postsynaptic elements of the Ia synapse
- Cathodal (hyperpolarising) tsDCS increases MN input resistance but does not impact Ia synapse morphology
- Both anodal and cathodal tsDCS fail to significantly modify the cellular burden of the disease

## Introduction

Amyotrophic Lateral Sclerosis (ALS) is a devastating neurological disease characterised by progressive loss of brainstem and spinal motoneurons (MNs). The burden of the disease is not shared equally between MN subtypes, as the MNs innervating fast fatigable muscle fibres (FFs) are preferentially targeted by the disease; MNs innervating fast resistant muscle fibres (FRs) degenerate later, and MNs innervating the slow muscle fibres (S) largely persist until the disease end-stage (Hegedus et al., 2008; Pun et al., 2006). These three types of spinal MNs display dramatically different activity patterns (Manuel and Zytnicki, 2011), and it has long been proposed that the cell-specific disease vulnerability is related to abnormal firing behaviour caused by an imbalance between cells’ intrinsic excitability and synaptic excitation levels (Pieri et al., 2003). According to this hyperexcitability theory, MN cell apoptosis is caused by excessive calcium influx during abnormally high cell firing rates driven by elevated glutamate levels (Ilieva et al., 2009; Van Den Bosch et al., 2006). However, clinical trials aimed at decreasing MN firing in the hope of slowing their degeneration have never produced clinically relevant results (Petrov et al., 2017); the only exception is Riluzole treatment, which prolongs patients’ survival in the terminal stage of the disease (Bensimon et al., 1994; Fang et al., 2018). On the other hand, chemogenetic interventions aimed at increasing the synaptic excitation of MNs have shown some positive effects on disease markers in animal ALS models (Bączyk et al., 2020a; Saxena et al., 2013). At this point, it is not clear if changes in excitability drive MN degeneration, or reflect homeostatic mechanisms aimed at maintaining the firing rates (Elbasiouny, 2022). However, it seems that disease-related changes in MN excitability follow a dynamic course (Huh et al., 2021). Interventions aimed at ameliorating the disease burden should therefore be targeted towards restoring MN excitability and excitation profiles to physiological levels, depending on the disease stage. However, pharmacological interventions aimed at altering MN electrophysiological profiles are hindered by a strong homeostatic response directed towards maintaining MN activity at the pre-programmed level (Joseph and Turrigiano, 2017). Indeed, numerous studies have shown neuronal desensitisation to pharmacological agents in the course of long-term administration (Antonucci et al., 2024; Löscher and Schmidt, 2006; Vinkers and Olivier, 2012).

It is therefore important to propose new ALS therapies that will utilise previously unexplored neuromodulation options. In this regard, the relatively new technique of trans-spinal direct current stimulation (tsDCS) appears an attractive candidate for ALS management. tsDCS is based on the polarisation of spinal circuits by externally applied electric fields and has already been shown to provide strong neuromodulation of both ascending and descending spinal tracts (Marangolo et al., 2020; Murray et al., 2018; Therkildsen et al., 2021; Yamaguchi et al., 2020) and also to spinal circuits (Lenoir et al., 2018; Pereira et al., 2022). Importantly, tsDCS effects are not diminished by desensitisation issues, as the MN electrophysiological profile is altered both by acute (Bączyk et al., 2020b, 2019) and chronic tsDCS application (Bączyk et al., 2020c).

In our recent article, we have shown that acute tsDCS application significantly and preferentially modifies the passive membrane properties and synaptic excitation levels of spinal MNs in the presymptomatic SOD1 G93A mouse model of the disease (Jankowiak et al., 2022). Building on these results, in this study we applied a two-week tsDCS protocol to presymptomatic (P30-P35) SOD1 G93A animals. We investigated the impact of the intervention on mouse spinal MN passive membrane properties, synaptic excitation levels, Ia synapse morphology, intracellular metabolic pathways activity, and disease markers. We hypothesised that repetitive alteration of MN excitability levels by tsDCS would provoke MN adaptation that could be detected at both electrophysiological and morphological levels. It will be shown that both anodal and cathodal tsDCS provoke strong neuroplasticity of SOD1 G93A mouse spinal MNs, with anodal polarisation acting towards increasing synaptic excitation and cathodal polarisation acting towards altering MNs’ passive membrane properties. These effects are linked to polarity-dependent alterations of Ia synapse morphology but have little effect on the intracellular pathway activity or the disease burden.

## Materials and Methods

### Animals

This study used a total of 48 male B6SJL-Tg (SOD1*G93A)1Gur/J mice (SOD1 mice henceforth) bred at the Wielkopolska Center of Advanced Technologies at the Adam Mickiewicz University (Poznań, Poland). Mice were housed two per cage at the Poznań University of Physical Education Animal Facility (Poznań, Poland) with unlimited access to food and water. The animal room was set to a reverse light/dark cycle (12 h/12 h) with humidity and temperature maintained at 55 ± 10% and 22 ± 2 °C, respectively. SOD1 mice display a phenotype similar to human ALS symptoms, with progressive limb muscle paralysis starting around postnatal day (P)90 and reaching end-stage ALS at around P120 (Gurney et al., 1994). To investigate the impact of tsDCS on the MN electrophysiological profile, 21 SOD1 mice were randomly assigned to a two-week tsDCS protocol, starting between P30 and P35; in vivo intracellular recordings were performed at P45–P55. Importantly, P45–P55 is considered a presymptomatic stage, just at the onset of fast-twitch muscle fibre denervation (Hegedus et al., 2008) when functional deficits are almost undetectable (Oliván et al., 2015) but MN pathology is evident (Delestrée et al., 2014; Martínez-Silva et al., 2018; Baczyk et al., 2020a). For immunohistochemical analysis, 27 SOD1 mice were subjected to tsDCS protocol, also starting at P30–P35. This cohort were sacrificed at P45–P55, and spinal cords were removed and processed. In accordance with European Union recommendations, our humane endpoints were defined as an inability of the mouse to reach food or water, the loss of more than 30% of body weight over 72 h, or an inability to rise or ambulate. None of the animals used in this study reached any of the humane endpoints. All procedures performed in this study were approved by the Poznań Local Ethical Committee (approval number 44/2018; Poznań, Poland). All authors held valid permits for working with laboratory animals and were appropriately trained in all experimental procedures.

### Two-week tsDCS to investigate MN synaptic excitation levels

tsDCS or sham polarisation was applied to SOD1 G93A animals randomly divided into Sham Control, Cathodal and Anodal polarisation groups. All groups underwent daily tsDCS or sham treatment lasting 15 minutes for a total of 14 days. For all groups, the tsDCS procedure started at P30–P35. To ensure adequate contact between the electrode and the skin, the mice were initially shaved on their back and abdomen. Each tsDCS session started with the administration of isoflurane anaesthesia (Isotek, Spain), initially in an animal induction chamber (Classical Vaporizer, VetFlo) using 4–5% isoflurane in air at a delivery rate of 100–200 ml/min for 5 minutes. Once a medium level of anaesthesia was reached (lack of hind limb withdrawal reflex, rhythmic breathing), the mice were moved from the induction chamber and the anaesthetic was delivered through a closely fitted nose cone. Thereafter, anaesthesia was sustained with 2–3% isoflurane in air (flow rate 100-200 ml/min). Application of a highly conductive electrolyte gel (Sigma Gel, Parker) to the animal’s skin preceded the placement of the electrode to enhance electrical conductivity and prevent skin damage. A rectangular stainless steel active electrode (5 × 10 mm) was placed above the Th13-L1 vertebrae; a metal crocodile clip attached to the abdominal skin flap ventral to the spinal electrode served as the reference electrode. Polarisation was applied using a custom-made battery-driven constant current stimulator (TP-1, WiNUE, Poland). By switching the polarity of the current source, this circuit arrangement can provide immediate current reversal from anodal to cathodal and vice versa. tsDCS was administered at an intensity of 100 µA, with either anodal or cathodal polarisation applied for 15 minutes, producing a current density of 2 µA/mm^2^ just below the active (dorsal) electrode. The current intensity was constantly monitored to ensure that the electrical charge delivered was the same for each animal. Once the polarisation was complete, the mice were moved to a cage to recover and were carefully observed for any signs of abnormal behaviour. The Sham Control group received an identical anaesthesia and electrode placement protocol; however, no current was passed through the electrodes.

### Two-week tsDCS to study the pre- and postsynaptic structures of the Ia-MN circuit

Sham Control, Cathodal and Anodal animals were subjected to a two-week tsDCS procedure identical to that described above. However, on day 10th of the protocol, shortly after the tsDCS / Sham session, the lateral gastrocnemius (LG) muscle was bilaterally injected with cholera toxin B subunit (CTB) to retrogradely label the LG MN pool. Following the injection, the tsDCS protocol was continued for the remaining 4 days. The CTB injection procedure is briefly described in the following paragraph, retrograde labelling of motoneurons.

### Retrograde labelling of motoneurons

The pool of LG MNs was retrogradely labelled in all mice subjected to the two-week tsDCS protocol (Sham Control, Cathodal and Anodal groups) with CTB conjugated with Alexa Fluor 555 (solution of 1.0 mg/ml CTB-555 with clean phosphate-buffered saline (PBS)). The animals were anaesthetised with isoflurane (2-3% in oxygen, as described in the above tsDCS protocol) and placed on a heating pad to maintain the central body temperature at 37 °C. A bilateral incision was made on the skin of the hindlimb just above the triceps surae (TS) muscle, and 5µl CTB-555 was injected into the LG muscle with a Hamilton microsyringe connected to a 22-gauge needle. Following the injection, the needle was removed, the skin was sutured with sterile ligatures and a subcutaneous injection of the analgesic and anti-inflammatory drug Meloxicam (0.4 mg/kg body weight) was given.

### Tissue processing for immunohistochemistry

The day after the final polarisation session (5 days after CTB injection), the animals were anaesthetised with a lethal dose of Morbital (200 mg pentobarbital/kg body weight, intraperitoneal). Mice were transcardially perfused (flow 6.5ml/min) with 40 ml 0.1M PBS and fixed with 70 ml ice-cold 4% paraformaldehyde (PFA). Isolated spinal cords were post-fixed in PFA overnight at 4°C and then cryoprotected in 30% sucrose in 0.1M PBS. The L3–L5 spinal cord segments were surrounded by Tissue-Tek® OCT cryo-embedding compound and frozen on dry ice. Transverse sections (30 μm) were cut on the cryostat (Thermo Scientific Microm HM 520) at -19°C and collected free-floating in cryoprotectant solution (300 ml glycerol, 500 ml 0.05M PB pH 7.4, supplemented with 150g sucrose) at -20°C.

### Immunostaining of pre- and postsynaptic structures

Free-floating sections containing CTB+ MNs (minimum 6 sections per animal) were washed in PBST (PBS + 0.2% Triton X-100) and incubated in a blocking buffer (5% normal donkey serum) in PBST for 2h at RT. Next, sections were incubated overnight at 4°C with goat anti-VGluT1 (vesicular glutamate transporter 1; 1:500, Synaptic Systems, cat. no 135 307) and rabbit anti-GluR4 (AMPA receptor subunit 4; 1:400, D41A11, Cell Signaling, cat. no 8070) antibodies diluted with PBST. Next, sections were washed in PBST, before 2h incubation at RT with the secondary antibodies (diluted at 1:500): donkey anti-goat CF-647 (Biotium 20048) and donkey anti-rabbit Alexa Fluor 488 (Thermo Fisher Scientific 21206). Afterwards, sections were rinsed three times and mounted in ProLong Gold Antifade (Thermo Fisher Scientific). Immunofluorescence (IF) labelling was carried out in one experimental session to ensure identical conditions of tissue processing and staining.

### Immunostaining of pCREB and disease marker

The free- floating sections with CTb+ MNs (minimum 5 sections per animal) were washed in PBST (PBS + 0.2% Triton X-100) and incubated in a 1% blocking buffer (ab126587) for 2h at RT. Next, sections were incubated 2-nights at 4°C with rabbit anti-phospho-CREB (Ser133) (1:200, Cell Signaling, mAb #9198), mouse anti-misfolded SOD1 (1:500, Mèdimabs, cat no MM-0070-P), guinea pig anti-VAChT (1:1000, Synaptic System, cat no 139 105) and goat anti-MMP9 (1:500, Sigma-Aldrich, cat no M9570) antibodies diluted with 1% blocking buffer in PBST. Next, sections were washed in PBST, prior to 2h incubation at RT with the secondary antibodies (diluted at 1:500): donkey anti-rabbit Alexa Fluor 488 (#711-545-152), donkey anti-mouse Alexa Fluor 647 (#715-605-150), donkey anti-guinea pig Alexa Fluor 405 (#706-475-148), donkey anti-goat Cy™3 AffiniPure™ (#705-165-003) respectively. Afterwards sections were rinsed 3 times and mounted in ProLong Gold Antifade.

### Image acquisition and quantification of pre- and postsynaptic structures

Fluorescent images were acquired with an Olympus FV1200 confocal inverted microscope with a 60x DIC oil-immersion objective (1.4 NA) and added 2.5 optical zoom. Twelve-bit depth images of CTB-labelled MNs, glutamate subunit 4 of the AMPA receptor (GluR4), and vesicular glutamate transporter 1 (VGluT1) consisting of digital slices were collected at 0.3 μm intervals with a pixel size of 0.082 μm. A minimum of 20 optical sections were acquired for each MN; MNs <30 μm in diameter or displaying less than two VGluT1+ synapses were excluded from the analysis. Images were collected at constant exposure parameters. Acquisition parameters were adjusted so as to maintain fluorescence intensity from target structures below oversaturation and above underexposure.

Image quantification was performed in ImageJ/Fiji (Schneider et al. 2012). For the evaluation of the number of presynaptic (VGluT1) and postsynaptic (GluR4) structures, seven consecutive optical sections were collapsed in a maximum-intensity projection, subjected to rolling-ball background subtraction. VGluT1+ boutons (objects larger than 1 μm^2^) located close to the cell body or proximal part of a dendrite (5 μm from the soma) and no further than 1 μm from the cytoplasm identified with CTB staining were taken to be analysed. Postsynaptic GluR4 structures colocalised with VGluT1, located between the presynaptic terminals and the cytoplasm of the cell were also analysed. The contour of the region of interest, including the area positive for VGluT1, was traced manually in ImageJ on created masks based on the IF signal of VGluT1 and automatically determined threshold values. The same method of analysis was used for GluR4 clusters. Any synaptic terminal opposing a cell body was analysed separately. The number, surface and mean fluorescence intensity of analysed structures were logged. The density of VGluT1 terminals was normalised to the cell surface determined by CTB labelling in the MN cytoplasm.

### Image acquisition and quantification of pCREB and misfolded SOD1 levels

Confocal images were obtained by using LSM710, Axio Imager 2 microscope (Zeiss, Jena, Germany) with a Plan Apochromoat 20x air objective with 0.8 NA. Images of motoneurons labeled with pCREB and misfSOD1 fluorescent probes were acquired using excitation wavelengths 488 nm and 647 nm. The imaging of motoneurons localized in L3-L5 lumbar region, labeled with VAChT and MMP9. Images were collected at constant parameters of exposure and all acquisition parameters were adjusted to maintain the emission fluorescence below oversaturation and above underexposure. All images were recorded as z-stack and merged using maximum intensity projection mode in ImageJ/Fiji software (National Institutes of Health). For the evaluation of pCREB and misfSOD1 immunofluorescent intensity, the contour of each nucleus and MN were traced manually on the maximum-intensity projection of confocal z-stack in ImageJ/Fiji and saved as ROI for quantification of the mean fluorescence intensity

### Preparation for electrophysiology

The methodology of electrophysiological experiments on spinal MNs in the mouse has been previously described in detail (Bączyk et al., 2020a, Jankowiak et al., 2022). Following premedication (atropine 0.20 mg/kg s.c., Polfa and methylprednisolone 0.05 mg, s.c., Pfizer), mice were anaesthetised by intraperitoneal injection of a drug cocktail (fentanyl 6.25 µg/ml (Polfa), midazolam 2.5 mg/ml (Polfa), medetomidine 0.125 mg/ml (Cp-Pharma), injected at 10 ml/kg body weight). A suitable depth of anaesthesia was determined by the lack of hind limb withdrawal reflex. Subcutaneous needle ECG electrodes were positioned for heart rate monitoring, and the central temperature was maintained at 37 °C with an infrared heating lamp and an electric blanket (TCAT-2DF, Physiotemp). After a tracheotomy was performed, the mouse was artificially ventilated with pure oxygen (SAR-1000 ventilator; CWE) with parameters adjusted to maintain the tidal CO_2_ level at around 4% (MicroCapstar; CWE). Right and left external jugular veins were catheterised for delivering additional doses of the anaesthetic cocktail (fentanyl 6.25 µg/ml (Polfa), midazolam 2.5 mg/ml (Polfa), medetomidine 0.125 mg/ml (Cp-Pharma); injected at 1.7 ml/kg body weight, i.v., administered every 20-30 min) or physiological buffer (4% glucose solution containing 1% NaHCO_3_ and 14% gelatine (Tetraspan; Braun) infused at 60 μl/h). The triceps nerve (innervating the TS muscle group, including medial gastrocnemius, LG, and soleus muscles) was dissected from the surrounding tissues. Two pairs of horizontal bars (Cunningham Spinal Adaptor; Stoelting) were used to immobilise the vertebral column between the T13 and L2, and a laminectomy was made at the T13–L1 vertebrae. The dura mater was removed from the exposed L3–L4 spinal segment to allow a glass microelectrode to be inserted into the spinal cord. The exposed tissues were covered with mineral oil. At the end of the surgery, animals were paralysed with pancuronium bromide (Pancuronium; Polfa; initial bolus of 0.1 mg, followed by additional doses of 0.01 mg every 30-40 min). From this point on, each additional dose of the anaesthetic was given at the same frequency as during the surgery, or when ECG and/or PCO_2_ levels neared the maximal physiological values.

### Stimulation and recording

MN electrophysiological properties were measured with glass micropipettes (tip diameter 1.0–1.5 μm, impedance 20-30 MΩ) filled with a mix of 2M K-acetate and 0.1M QX-314 (sodium channel blocker; Sigma-Aldrich). Intracellular recordings were obtained with an Axoclamp 900A amplifier (Molecular Devices) connected to a Power1401 interface (CED, sampling rate 20kHz) operated by Spike2 software (CED). The amplifier system was used in bridge mode to record excitatory postsynaptic potentials (EPSPs; single response or paired-pulse responses) or in discontinuous current clamp (DCC) mode (switching rate 7-8 kHz) to record the responses to square pulses necessary to determine input resistance. Peripheral stimulation of the triceps nerve enabled us to identify MNs based on an “all-or-nothing” antidromic action potential. This method of MN antidromic identification was still possible since it takes 30–90 s for the QX-314 to diffuse from the electrode tip to the cell body and block the antidromic spike. The TS nerve was stimulated with constant current pulses of 0.1 ms duration and amplitude up to 50 µA delivered at 3 Hz (DS4, Digitimer), but not more than 2.5 x threshold (Th) of the most excitable fibres in the nerve. An electrode positioned at the dorsal surface of the spinal cord was used to record the group I afferent volley in response to peripheral nerve stimulation (Fig.1A_1_). All the MNs selected for the analysis were characterised by a resting membrane potential (RMP) more hyperpolarised than -50 mV and an initial overshooting action potential. The maximal EPSP amplitudes were measured from the RMP (measured just after the stimulation artefact) to the most depolarised part of the voltage response (Fig.1A_2_) and EPSP time to peak was measured from the beginning of EPSP to the maximal amplitude. The EPSP half decay time constant was measured by plotting an exponential curve to the descending part of the EPSP trace, to the point when it decayed 50% of the maximal EPSP amplitude (Fig.1A_2_). In the paired-pulse (PP) protocol, the peripheral nerve was stimulated with a pair of identical constant-current stimuli of 0.1 ms duration separated by 10 ms intervals that elicited EPSPs (Fig. 2C). The PP ratio was calculated by measuring the amplitude of the second EPSP measured from the most hyperpolarised part of the voltage trace after the first EPSP from the pair, and dividing it by the amplitude of the first EPSP measured from the RMP. This method of PP analysis is independent of possible temporal summation of the EPSP traces, which can occur if the EPSP decay is especially long (Jackman et al., 2016; Jiang and Abrams, 1998; Zhang and Schneider, 2011). The PP ratio was measured for each individual EPSP pair and then averaged across the recording period (at least 10 EPSP pairs per average). The MN input resistance at RMP (RIN) was calculated by measuring the membrane voltage deflection in response to a series of small-amplitude square current pulses (-2 to +2 nA, 500 ms), as described previously (Manuel et al., 2009, and outlined in Fig. 1A_3_). Injecting small hyperpolarising current pulses (-5 nA, 1 ms) enabled us to measure the membrane time constant on the relaxation of the membrane potential (TauM). At the end of the experiment, animals were euthanised with a lethal intravenous dose of sodium pentobarbital (200 mg/kg).

**Figure 1.**
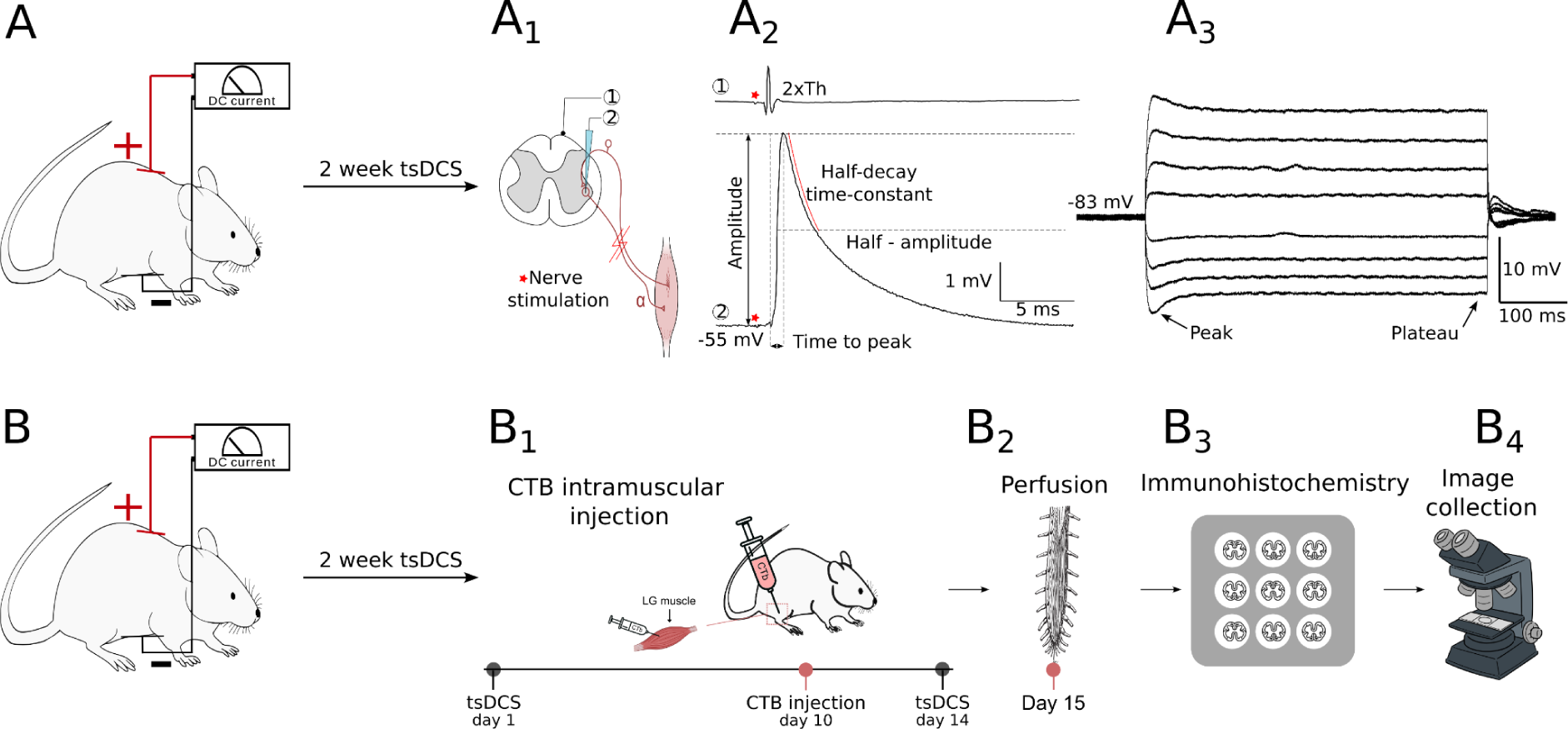
Experiment design. **A**, P30–P35 SOD1 G93A mice were subjected to two weeks of Sham Control, Cathodal or Anodal polarisation protocol. One day after the last polarisation session, an electrophysiological experiment was performed as outlined in **A_1_** and the Ia EPSPs (**A_2_**) and passive membrane properties (**A_3_**) of spinal motoneurons (MNs) were investigated. **B**, as in **A**, P30–P35 mice were subjected to two weeks of Sham Control, Cathodal or Anodal polarisation; however, at the 10^th^ day of polarisation protocol (**B_1_**), a bilateral injection of cholera toxin B (CTB) was made into lateral gastrocnemius (LG) muscles of all animals to retrogradely label the LG MN pools. **B_2_ – B_4_**, one day after the last polarisation session, mice were sacrificed, spinal cords removed and immunohistochemical analysis of MN synaptic coverage, intracellular metabolic pathways activity and misfolded SOD1 protein levels was performed.

**Figure 2.**
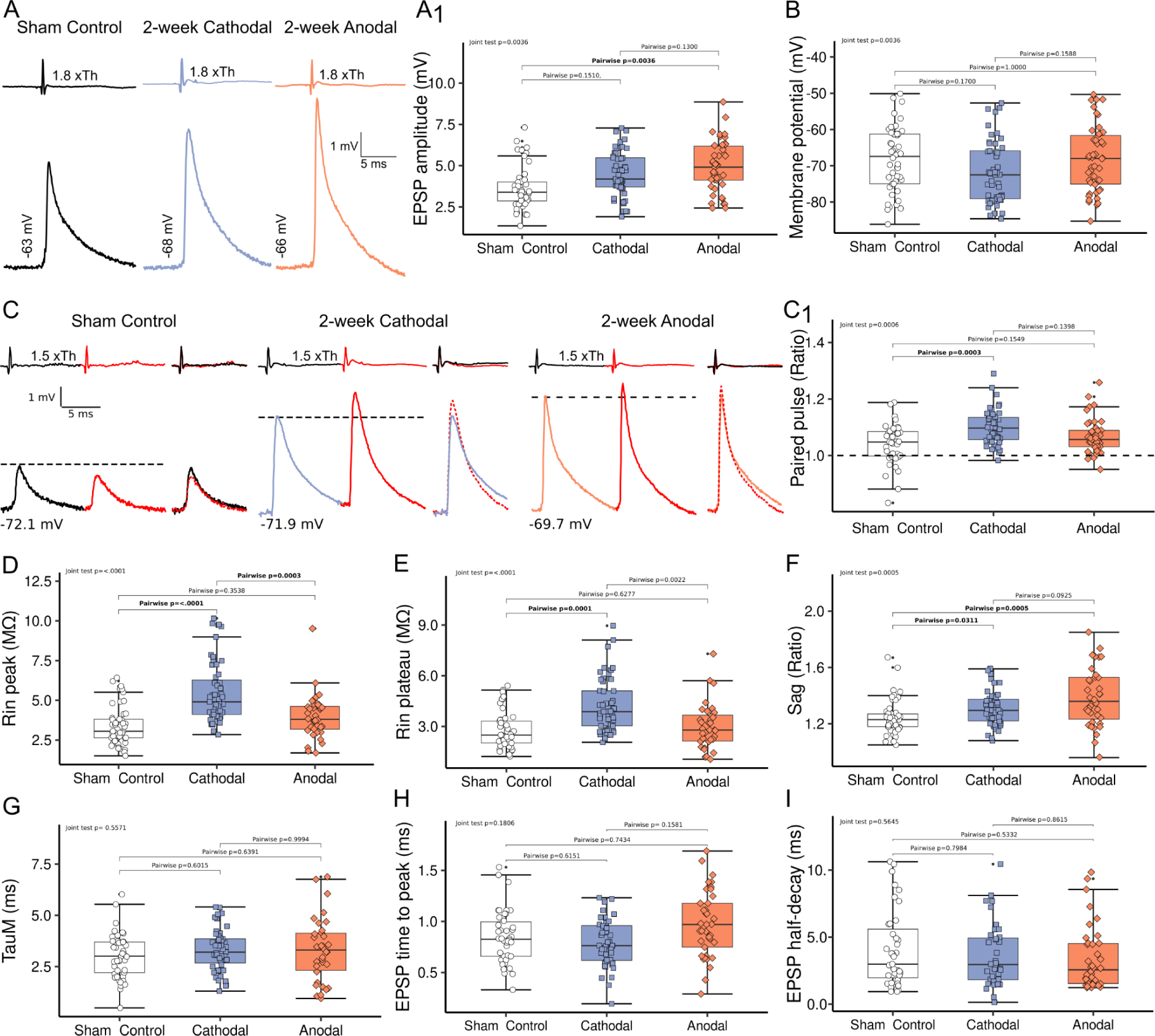
The effects of two-week polarisation protocol on SOD1 G93A mice motoneuron electrophysiological profile. **A**, intracellular records of maximal Ia EPSPs recorded from triceps surae (TS) motoneurons (MNs) of Sham Control, Cathodal or Anodal polarisation groups. The upper trace shows a group I volley recorded from the surface of the spinal cord evoked by TS nerve electrical stimulation. The 1.8xTh (1.8xThreshold) on the upper trace corresponds to the intensity of TS nerve stimulation. The lower trace shows maximal Ia EPSPs recorded from TS MNs evoked by TS nerve stimulation. Notice a significantly higher maximal EPSP amplitude for the Anodal polarisation group. **A_1_**, **B,** box plots showing the data distribution from Sham Control, Cathodal and Anodal polarisation groups for maximal Ia EPSPs and resting membrane potential respectively. **C**, intracellular recordings of MNs subjected to the paired-pulse protocol from Sham Control, Cathodal and Anodal polarisation groups. Panel description as in **A**. For each test, the second response is marked red and superimposed as a dotted line on the first response to the right of the original trace. **A**, dashed line indicates the 1st response amplitude. Notice a paired-pulse depression in a large subset of MNs in the Sham Control group. This phenomenon is largely reduced in the Cathodal polarisation group with only a single MN showing paired-pulse depression. **C_1_**, box plot description as in **A_1_** and **B**, but showing the paired-pulse protocol results with the dashed line showing the border between paired-pulse depression and paired-pulse facilitation. **D–F**, box plots showing data distribution of peak (**D**) and plateau (**E**) input resistance, SAG ratio (**F**), membrane time constant (**G)**, EPSP time to peak (**H**) and EPSP half-decay time constant (**I**). Each point on the boxplot indicates a single MN. The lower and upper hinges correspond to the first and third quartiles (25th and 75th percentiles, respectively). The upper whisker extends from the hinge to the largest value no further than 1.5x the interquartile range (IQR). The lower whisker extends from the hinge to the smallest value, at most 1.5x the IQR. Outliers were checked for correctness and are plotted individually if they meet all inclusion criteria. The significant differences between the investigated groups are marked in bold at **p<0.05**.

### Statistics

All plots and statistical analyses were programmed in RStudio 2023.06.1 (Posit Software, PBC) with appropriate libraries. Plots were created with the ggplot2 package (Hadley Wickham, 2016). A generalised linear mixed model, including zero-inflation and dispersion corrections, was plotted using the glmmTMB package (Brooks et al., 2017) to statistically compare electrophysiological and immunohistochemical data, and to account for the effect of individual differences on MN properties (Highlander and Elbasiouny, 2022). The fixed effect factor was the polarisation (Sham Control, Cathodal or Anodal), while Mouse was the random effect. To validate the fitted model, a Residual Diagnostics for HierArchical (Multi-level / Mixed) Regression Model was performed using the DHARMa package. A plot of the scaled residuals was created by simulating from the fitted model, and data deviation, dispersion and variance were assessed. If a significant deviation from the model was found, then the dispersion correction or data family correction was introduced and the model was refitted. Only the models that met all assumptions were used for further analysis. The significance of the fixed effect was established with joint tests of the terms in a model, and between-group significance was then assessed with estimated marginal means for specified factors from the emmeans package (Russell V. Lenth, 2023), with the Tukey adjustment method for comparing a family of three estimates. The significance level of the fixed effect was set at p<0.05. For each comparison, we report the Estimated Marginal Mean value ± SE. To estimate the correlation of EPSP properties with MN passive membrane properties, the Spearman correlation coefficients were analysed using the ggpubr package (v0.6.0 Kassambara A, 2022) and the p values are given with significance level set at p<0.05. In the box plots, each data point represents a single MN for electrophysiological data, and a single synapse or a single MN for immunohistochemical data.

## Results

### Two weeks of anodal tsDCS significantly increases EPSP amplitudes in SOD1 mice

In the first series of experiments, we investigated whether two-week tsDCS triggers MN plasticity, measured as a change in the levels of MN synaptic excitation by Ia afferents. The analysis was performed on n=47 MNs in the Sham Control group (n=5 mice), n=52 MNs in the Cathodal group (n=10 mice) and n=42 MNs in the Anodal polarisation group (n=6 mice). Figure 2 shows that anodal polarisation repeated 15 min daily for two weeks significantly increased the maximal Ia EPSP amplitudes recorded from TS MNs of SOD1 G93A mice. The maximal amplitudes of unitary Ia EPSPs were on average 38% larger in the Anodal polarisation group when compared to the Sham Control group (5.21±0.34 mV vs. 3.77±0.25 mV, Joint test F(2, 136)=5.866, p=0.0036, post-hoc vs. Sham Control Tukey p=0.0028, Fig. 2A, A_1_). In contrast, no change in maximal EPSP amplitude was seen in the Cathodal polarisation group (post-hoc p=0.1510, Fig. 2A, A_1_). The increase of the EPSP amplitude by anodal polarisation was not followed by any changes in the MNs’ passive membrane properties (Fig. 2B, D-G, all tests p>0.05). However, a strong increase of both peak (5.44±0.236 MΩ vs. 3.35±0.268 MΩ, Joint test F(2, 128)=18.596, p<0.0001, post-hoc vs. Sham Control, Tukey p<0.0001, Fig. 2D) and plateau input resistance (4.21±0.224 MΩ vs. 2.70±0.204 MΩ, Joint test F(2, 128)=13.246, p<0.001, post-hoc vs. Sham Control Tukey p=0.0001, Fig. 2E) was seen in the Cathodal polarisation group. Furthermore, a significant increase in the SAG ratio was seen in both Anodal and Cathodal polarisation groups (1.41±0.0378 and 1.31±0.0219 vs. 1.23±0.0247 for Andoal and Cathodal vs. Sham Control groups respectively, Joint test F(2, 123)=8.144, p=0.0005, post-hoc vs. Sham Control Tukey p=0.0005, and p=0.0311). Interestingly, further analysis showed that cathodal polarisation significantly increased the PP ratio (1.10±0.0115 vs. 1.04±0.0146 Joint test F(2, 108)=4.983, p=0.0085, post-hoc vs. Sham Control, Tukey p=0.0003, Fig. 2C, C_1_). This is in sharp contrast to the multiple cases of pathological PP depression seen in the Sham Control group, as we have previously reported for SOD1 animals (Bączyk et al., 2020a) (Fig. 2C, C_1_).

Next, we analysed the time course of EPSPs to determine possible alterations of AMPA receptor dynamics and Ib inhibition. While Ib inhibition from tendon organs can appear at a stimulation intensity close to the Ia afferents activation threshold (Eccles et al., 1957; Jankowska et al., 1981), it does not affect the maximal amplitude of Ia EPSPs in mouse MNs, as the maximal EPSP rise time is faster than the latency of disynaptic inhibition (Bączyk et al., 2020a). Still, alterations of Ib inhibition should be detectable as changes in the EPSP half-decay time constant. Both maximal EPSP rise time and the maximal EPSP half-decay time constant were not significantly different between the studied groups (Joint test F(2, 100)=0.484, p=0.6175 and Joint test F(2, 100)=0.998, p=0.3724 for EPSP half-decay time and EPSP rise time respectively, Fig. 2H, I) indicating that the possible Ib inhibition component of the EPSP response was not significantly different between the analysed groups.

Previous data has shown that in WT rats, Ia EPSP amplitudes are strongly correlated with cell RMP and peak input resistance (Krutki et al., 2023, 2022). Since in this study in SOD1 mice, the peak input resistance was significantly increased in the Cathodal polarisation group (Fig. 2D), we set out to determine if these changes could explain the observed EPSP alterations. However, there was very little correlation between EPSP amplitude and peak input resistance in all analysed groups (Spearman, R=-0.04, p=0.79; R=-0.066 p=0.65; and R=-0.067, p=0.72; for Sham Control, Cathodal and Anodal polarisation groups, respectively) indicating that RIN variations are not responsible for the change of EPSP amplitude after tsDCS (Fig. 3A). On the other hand, as in the previous rat studies (Krutki et al., 2022) a weak negative correlation was seen in our study between EPSP amplitude and MN RMP, though it was significant only in the Cathodal polarisation group (Spearman, r=-0.079, p=0.6; R=-0.39, p=0.0046; and R=-0.28, p=0.082; for Sham Control, Cathodal and Anodal polarisation groups, respectively, Fig. 3B). Further, a significant negative correlation was present in the Anodal polarisation group between the maximal EPSP amplitude and EPSP time to peak (Spearman, R=-0.53, p=0.00071) and between the maximal EPSP amplitude and EPSP half-decay time (Spearman, R=-0.37, p=0.021). For the Sham Control and Cathodal polarisation groups, the correlations between the maximal EPSP amplitude and EPSP time course were not significant (All Spearman p>0.5)

**Figure 3.**
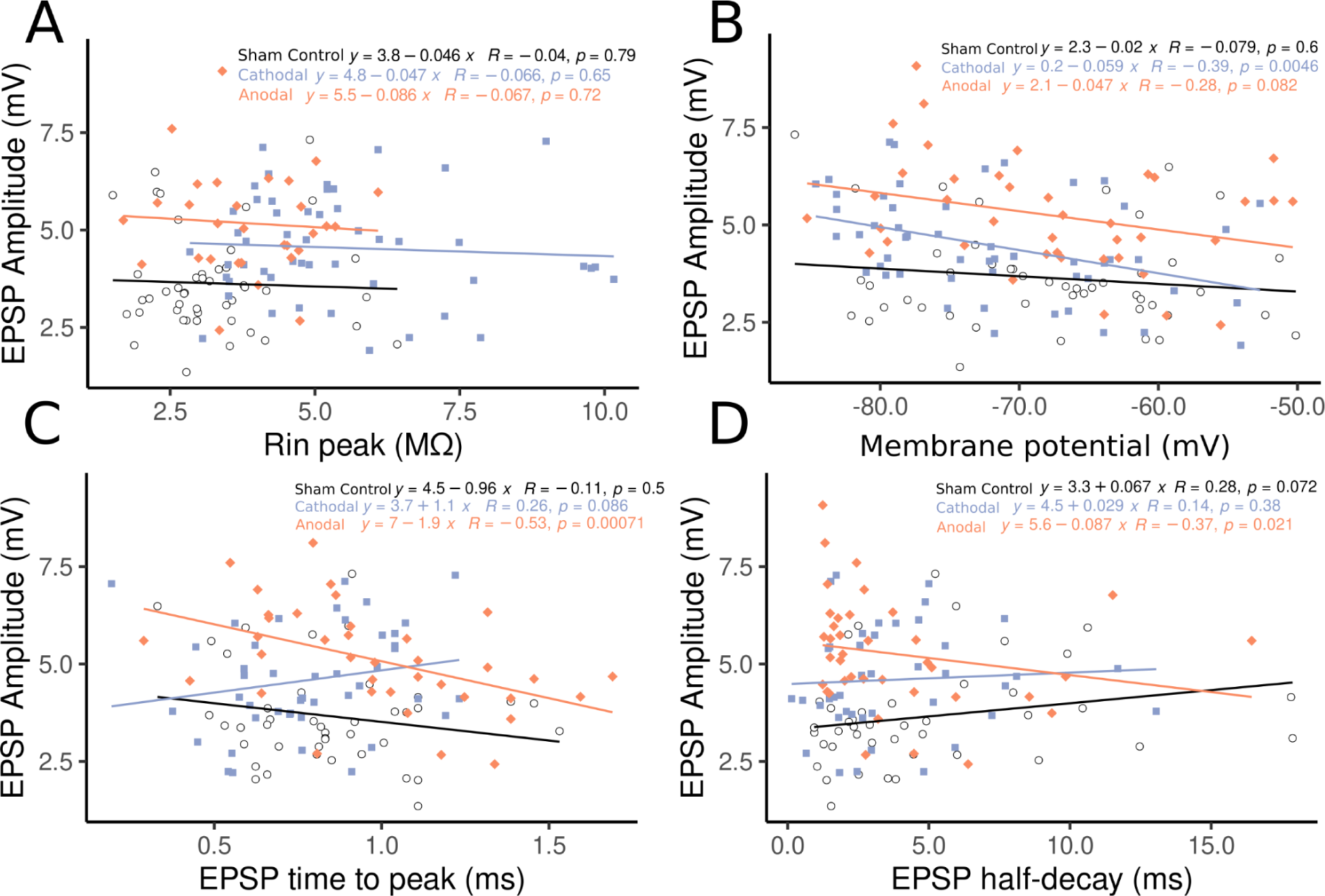
Correlation of maximal EPSP amplitudes with MN passive membrane properties and EPSP time course. Maximal EPSP amplitudes from Sham Control, Cathodal and Anodal polarisation groups are plotted against peak input resistance **(A)**, resting membrane potential **(B)**, EPSP time to peak **(C)** and EPSP half-decay time constant **(D)**. For each relationship, a linear trendline is plotted and the Spearman correlation coefficient (R) with p-value is shown on the right of the trendline equation. On the plots, each data point represents a single MN.

Taken together these results show that two-week tsDCS evokes strong neuronal plasticity in SOD1 mice, with anodal polarisation acting towards increased synaptic excitation levels.

### Two weeks of tsDCS significantly affects Ia synapse morphology

Having identified the impact of tsDCS on MNs’ electrophysiological profile and synaptic excitation levels, we set out to determine if the observed changes can be linked to morphological changes in the Ia synapse. We first quantified the level of fluorescence intensity of Ia synaptic boutons, identified by the presence of VGluT1 on the soma of CTB-identified LG MNs in spinal cords harvested from mice subjected to the Sham Control (n=5), Cathodal (n=5) or Anodal (n=5) polarisation protocol. Importantly, while all proprioceptive afferents express VGluT1 (and not VGluT2), only the Ia projects to spinal MNs, and therefore the VGluT1 boutons on MNs can be ascribed to Ia terminals. Figure 4 shows a strong polarity-dependent plasticity of Ia monosynaptic terminals in response to two types of polarisation. The level of fluorescence intensity of VGluT1 boutons on LG MNs was not significantly different between either polarisation and Sham Control groups (Joint test, F(2, inf)=4.592, p=0.0101, post-hoc vs. Sham Control p=0.1236, and p=0.6922 for Anodal and Cathodal polarisation groups, respectively, Fig. 4A, C, G). However, a strong difference was seen in the VGluT1 IF between the Anodal and Cathodal polarisation groups, with the Anodal group displaying a 13.5% stronger IF signal (1621±50.6 AU vs. 1412±49.0 AU, Tukey p=0.0084). A similar effect was seen for the average VGluT1 bouton size, with no significant difference when either group was compared to Sham Control (Joint test, F(2, inf)=0.0183, post-hoc vs. Sham Control Tukey p=0.5893 and p=0.2060 for Anodal and Cathodal polarisation, respectively), and a significant 14% larger bouton size in the Anodal group when compared to the Cathodal group (2.91±0.147 µm^2^ vs. 2.35±0.132 µm^2^, Tukey p=0.0136, Fig. 4H). However, the density of VGluT1+ synapses on the MN membrane was not affected by either polarisation protocol (Joint test F(2, inf)=0.968, p=0.3797, Fig. S1 B), indicating that the difference of VGluT1 fluorescence signal between polarisation groups is not caused by synaptic sprouting.

**Figure 4.**
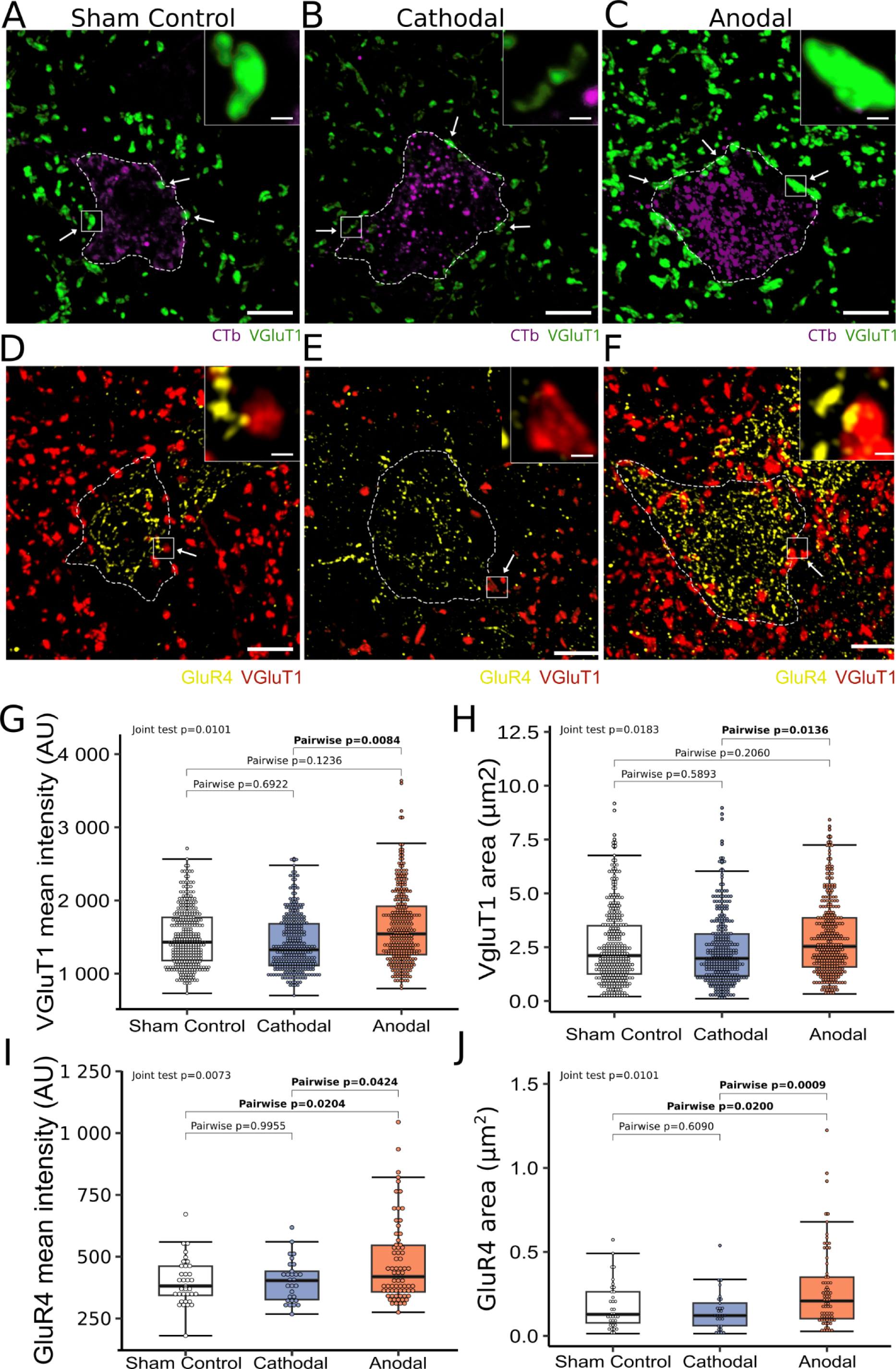
VGluT1 and GluR4 response to two weeks tsDCS. **A–C,** tsDCS-dependent changes in VGluT1 fluorescence intensity (AU) and area (μm^2^). Each large panel depicts a single motoneuron (MN) (white dashed outlines indicating the approximate cell body shape, scale bar: 10 μm) with a magnified region illustrating VGluT1 synapse contact on the MN cell body (scale bar: 1 μm). White arrows mark VGluT1 synaptic contacts on the MN surface. **D,** the box plot displays the distribution of VGluT1 fluorescence intensity (AU) across three experimental conditions: Sham Control, Cathodal, and Anodal polarisation groups, respectively. Notice a significantly higher VGluT1 fluorescence intensity in the Anodal compared to the Cathodal polarisation group. **E,** The box plot illustrates alterations in VGluT1 bouton area (μm^2^) across experimental conditions. **F–H,** each large panel depicts a single MN (white dashed outlines indicating the approximate cell body shape, scale bar: 10 μm). Within each panel, a magnified square region illustrates the post-synaptic GluR4 cluster juxtaposed to the presynaptic VGluT1 bouton (scale bar: 1 μm). **I,** the box plot displays the distribution of GluR4 fluorescence (AU) signal across three experimental conditions: Sham Control, Cathodal, and Anodal polarisation, respectively. A significantly stronger IF signal can be seen in the Anodal polarisation group when compared to both the Cathodal and Sham Control groups. **J**, the box plot illustrates the distribution of the GluR4 cluster area at VGluT1 synapses across three experimental conditions: Sham Control, Cathodal, and Anodal polarisation groups, respectively. Again, the GluR4 cluster area is significantly larger in the Anodal than in the Cathodal and Sham Control groups. Boxplots and statistical description as in Fig. 2.

Having identified changes in the VGluT1 contacts, we then asked if the postsynaptic elements of the Ia synapse are also affected by tsDCS. To this end, we quantified the level of fluorescence intensity of the GluR4 subunit of the AMPA receptor, as our previous investigations linked their lower expression to reduced maximal EPSP amplitudes in SOD1 animals (Bączyk et al., 2020a). Importantly, GluR4 is recognised as the principal post-synaptic subunit found in MNs at VGluT1 synapses (Ragnarson et al., 2003). We were able to detect a significant increase in the GluR4 IF signal intensity following the Anodal polarisation protocol (461±14.4 AU vs. 400±17.4 AU, Joint test F(2, inf)=4.926, p=0.0073, post-hoc vs. Sham Control Tukey p=0.0204, Fig. 4D-F, I); no change was observed following the Cathodal polarisation protocol (post-hoc vs. Sham Control p=0.9955, Fig. 4I). Further analysis showed that the area of the GluR4 was also significantly larger in the Anodal than in the Sham Control group (0.280±0.0267 vs. 0.180±0.0243 µm2, Joint test F(2, inf)=7.808, p=0.0004, post-hoc vs. Sham Control Tukey p=0.0280), while no significant effect of the Cathodal polarisation on the GluR4 cluster area was detected (post-hoc vs. Sham Control Tukey p=0.6090, Fig. 4J). These results consistently show that the polarisation protocol triggered strong plasticity of the Ia synapse, expressed predominantly in the recovery of GluR4 subunits of the AMPA receptor.

### Two weeks of tsDCS does not change intracellular pathways activity and disease marker levels

Our previous work showed that a chronic increase of MN synaptic excitation levels induced by chemogenetics ameliorates disease burden, as indicated by a reduction of misfolded SOD1 protein levels (Bączyk et al., 2020a). In these experiments, a similar, strong increase of MN synaptic excitation was observed following two weeks of anodal polarisation, and in parallel, a strong increase in input resistance was seen following cathodal polarisation (Fig. 2). Both of these effects may translate into stronger MN activity (Manuel and Zytnicki, 2011), which in turn can activate intracellular pathways and provide activity-dependent neuroprotection (Bączyk et al., 2020a). Therefore, in the final set of experiments, we set out to investigate whether the observed tsDCS effects translate into changes in intracellular pathway activity and disease marker levels. In opposition to our expectations, tsDCS had a negligible effect on the analysed parameters. Figure 5 shows that indeed, both cathodal and anodal polarisation failed to produce a significant change in the levels of phosphorylated pCREB, a marker of activity-dependent intracellular pathways activity (Joint test F(2, inf)=0.8281, all post-hoc p>0.05, Fig. 5A-C, D). Similarly, no significant effect was seen on the levels of an intracellular disease marker—the misfolded SOD1 protein (Joint test F(2, inf)=0.6031, all post-hoc p>0.05, Fig 5A-C, E).

**Figure 5.**
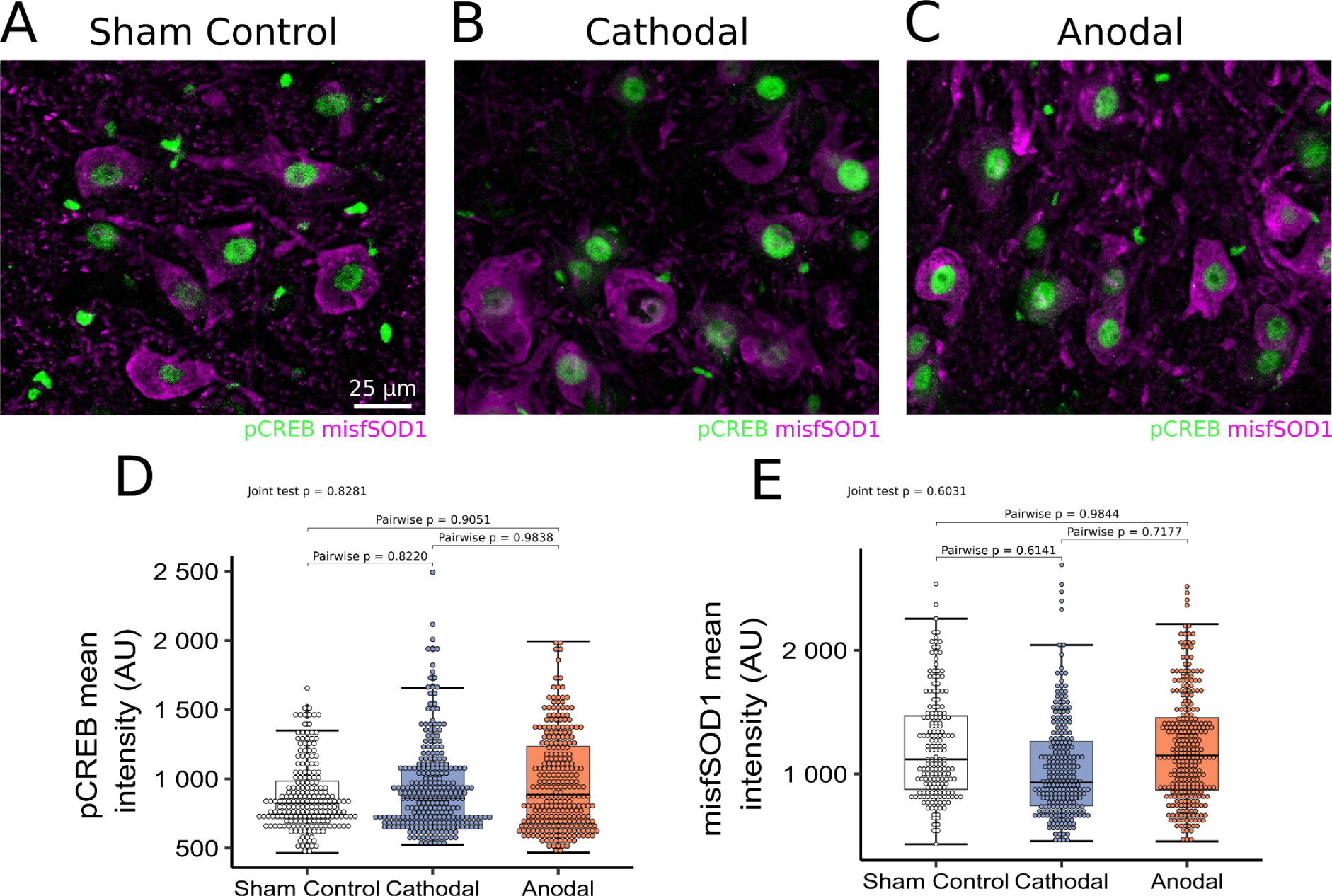
Modest tsDCS effects on pCREB and misfSOD1 levels. **A-C,** tsDCS-dependent changes in pCREB and misfolded SOD1 (misfSOD1) fluorescence intensity (AU) in L3–L4 motoneurons. **D,** the box plot displays the distribution of pCREB fluorescence intensity (AU) across three experimental conditions: Sham Control, Cathodal, and Anodal polarisation groups, respectively. **E,** as for **D**, but displaying data distribution for misfSOD1 fluorescence intensity (AU). Note a lack of significant effect on both pCREB and misfSOD1 levels. Boxplot and statistical description as in Fig. 4.

## Discussion

This study is the first to investigate the impact of a two-week tsDCS protocol applied to SOD1 mice at a pre-symptomatic stage of modelled ALS. We show a strong polarity-dependent effect of tsDCS on the electrophysiological profile and Ia synapse morphology of SOD1 mouse MNs. The effects of anodal tsDCS are predominantly facilitatory and include an increase of maximal Ia EPSP amplitude, an increase in VgluT1 fluorescence intensity and restoration of the GluR4 subunit of the AMPA receptor. On the other hand, the effects of cathodal polarisation are modest in their effect on EPSP amplitude but show a strong increase in MN input resistance and recovery of the PP ratio. While the effects of polarisation on the spinal MN electrophysiological profile and Ia synapse morphology are apparent, they do not translate into significant alterations of the intracellular tract activity and they do not modify the disease burden.

### Anodal tsDCS increases EPSP amplitude with no change in passive properties

The average amplitude of maximal EPSPs in the Anodal polarisation group (5.28±0.33 mV), was indeed very similar to maximal Ia EPSPs in untreated WT mice (4.98±1.57 mV), as presented in (Jankowiak et al., 2022). This indicates that our polarisation protocol fully recovered the disrupted maximal Ia EPSPs of SOD1 animals to physiological values. This recovery may be caused by either cell-autonomous mechanisms acting on the cell’s membrane properties, such as input resistance (Manuel and Zytnicki, 2011), RMP (Eccles et al., 1962; Edwards et al., 1976), activity of several types of inward and outward ion currents (George et al., 2009; Sharples and Miles, 2021), and number of available AMPA receptors at the Ia synapse (Burke et al., 1988; Montes et al., 2015); or by extracellular factors such as an increase Ia synapse number and density (Lev-Tov et al., 1983), or the levels of available neurotransmitter at presynaptic release sites. Due to our experimental setup utilising electrical stimulation of Ia afferents, we can exclude a possible contribution of altered muscle spindle activation that could otherwise impact the EPSP amplitude. Our results indicate that it is also unlikely that the increase of maximal Ia EPSP amplitude in the Anodal polarisation group was caused by the alteration of passive membrane properties, as the cell’s membrane potential, time constant and input resistance were not significantly affected. This is further confirmed by a lack of increase in maximal EPSP amplitude in the Cathodal polarisation group, in which we found a strong increase in the peak input resistance.

On the other hand, an increase in the SAG ratio was found in both the Anodal and Cathodal polarisation groups, indicating elevated activity of the hyperpolarisation-activated Ih current (Sharples and Miles, 2021). The activation of the Ih current can reduce the EPSP amplitude (George et al., 2009), but we did not see this in our data sample. This might be explained by two different compensatory mechanisms. In the Anodal polarisation group, a possible decrease of maximal EPSP amplitude by active Ih could have been counteracted by increased VgluT1 levels and recovery of the GluR4 subunit of the AMPA receptor (see discussion below). In the Cathodal group, the increase in peak RIN allowed more effective EPSP generation (Manuel and Zytnicki, 2011) and prevented the Ih current-related EPSP decrease.

It therefore seems that alterations in MN passive membrane properties play only a minor role in the tsDCS-induced alterations of EPSP amplitudes.

### Low correlation between MN passive membrane properties and maximal Ia EPSP amplitudes may be a unique feature of mouse MNs

In our study, the MN peak input resistance showed very little correlation with the maximal amplitude of Ia EPSPs. This is highly surprising, as a strong correlation between these parameters is apparent in both rat (Krutki et al., 2022) and cat (Heckman and Binder, 1988) MNs. The intracellular recordings of maximal Ia EPSPs in mouse MNs have so far only been reported in a single paper (Jankowiak et al., 2022); it is therefore difficult to judge whether a lack of correlation between the peak RIN and maximal EPSP amplitude is a consequence of the SOD1 mutation in our animals, a result of the polarisation protocol, or a general phenomenon of all mouse MNs. Indeed, the latter hypothesis is plausible, as a re-analysis of data from (Jankowiak et al., 2022) showed no correlation between peak RIN and maximal Ia EPSP amplitude in both WT and SOD1 mice in control conditions (Fig. S2). At this point, we are unable to explain this phenomenon, however, several lines of evidence indicate that indeed, the mouse MNs have some properties not encountered in other species. First, in mice, the majority of motor unit force is produced at a sub-primary range of MN firing (Manuel and Heckman, 2011) and not in the primary range, as in the case of rat (Manuel et al., 2019) or cat motor units (Burke, 1968). Second, mouse and cat MNs differ in the amplitude of the persistent inward current, which does not scale with the MN size (Huh et al., 2017) and allows mouse MNs to generate action potentials at significantly higher frequency. Third, mouse MNs do not exhibit a secondary range of firing in response to triangular ramp injection (Manuel et al., 2009), which is again observed in both rat (MacDonell et al., 2012) and cat (Kernell, 1965) MNs. Therefore, it is possible that mouse MNs do not possess the classical EPSP amplitude/peak input resistance relationship seen in other species.

### Anodal and Cathodal tsDCS differentially affect Ia synapse dynamics

Significant negative correlations between the maximal EPSP amplitude, the EPSP time to peak, and the EPSP half-decay time constant were seen in SOD1 mice MNs following the anodal polarisation protocol. This shows that the faster EPSP dynamics were related to more efficient EPSP generation. On the other hand, this relationship (although slightly below the level of significance) was reversed in the MNs subjected to the cathodal polarisation protocol. While the average EPSP rise time and half-decay time constants were not different between the analysed groups (Fig. 2), these results indicate that anodal and cathodal tsDCS have differential effects on Ia synapse dynamics.

### Anodal tsDCS promotes activity-dependent plasticity at VGlut1 synapses

The intensity of the VGlut1 signal was significantly increased in LG MNs of SOD1 mice following two weeks of anodal tsDCS. This indicates that a greater quantity of neurotransmitter was available in the synaptic vehicles at the presynaptic side of the Ia synapse; together with the recovery of the AMPA GLuR4 receptor subunit, this could explain the increased maximal Ia EPSP amplitudes recorded in the MNs of SOD1 animals. The increased neurotransmitter levels are most likely related to the increased excitability of the Ia afferent fibres, resulting in stronger synaptic input to spinal MNs. In our previous studies, we have shown that a single 15-minute session of anodal tsDCS provokes long-term increases in Ia afferent activity (Jankowiak et al., 2022). Similarly, the works of Jankowska and Hammar (Jankowska and Hammar, 2021) show that tsDCS-induced afferent neuromodulation can last up to 2 hours following treatment. Therefore, it might be expected that the excitability of Ia afferents would increase in tsDCS-treated SOD1 mice and that this increase would last long after the end of the tsDCS session. When the mice recover from anaesthesia following a tsDCS session and return to normal activity, the prolonged increase in the Ia fibre excitability translates into elevated Ia afferent input to spinal MNs from contracting muscle fibres. This might trigger adaptive changes in Ia synapses on spinal MNs, similar to after-training protocols involving strong Ia afferent activation (Krutki et al., 2022). A similar phenomenon was reported by Gajewska-Woźniak et al. (Gajewska-Woźniak et al., 2016), who found an increased fluorescence signal of glutaminergic terminals on MNs following two weeks of low-threshold afferent stimulation, which preferentially activated Ia and Ib fibres. Furthermore tsDCS was recently found to increase MN firing by increasing calcium influx by acting on L-type calcium channels (Song and Martin, 2022). This can result in an increased activation of the intracellular metabolic pathways such as the cAMP-dependent protein kinase (PKA) pathway, which would further act on the phosphorylation of AMPA receptors (Esteban et al., 2003) and provoke synaptic plasticity. In our previous works, we have already shown that chronic PKA activation recovers the deranged post-synaptic elements of the Ia synapse and partially rescues the Ia EPSPs amplitude (Bączyk et al., 2020a). Since this investigation shows both a recovery of Ia EPSP amplitude and an increase in the levels of GluR4 subunits of the AMPA receptors in the anodal-treated animals, we believe that an increase of PKA activity is one of possible mechanism of the polarity-induced alterations. However, the lack of increase of pCREB levels indicates that this was not sufficient to restore MN physiological activity levels.

### Changes in the paired-pulse ratio point to differential effects of anodal and cathodal polarisation

We found that the PP ratio was significantly increased by cathodal polarisation, while anodal polarisation had only a modest effect on this parameter. This might be controversial given the fact that anodal polarisation increased EPSP amplitudes and largely recovered AMPA GluR4 subunits. However, this result is in line with our previous data showing a decrease in the PP ratio following a single session of anodal polarisation, and no changes following a single session of cathodal polarisation (Fig. S3). Similarly, the recent work of Nascimento et al. (Nascimento et al., 2024) shows that an increase in the quantal release at the Ia synapse in neonatal SOD1 animals leads to an increase in spinal MN synaptic excitation and a concurrent reduction of the PP ratio. The principle of the PP facilitation phenomenon is the release of the tonic presynaptic inhibition affecting the primary afferents following the first shock of the stimulation doublet; this allows more neurotransmitters to be released into the synaptic cleft making the second EPSP larger (Stuart and Redman, 1991). However, the amplitude of the EPSP does not directly depend on the amount of released neurotransmitter but is also connected to the number of activated postsynaptic receptors. We have already shown that in SOD1 mice, the MN PP ratio is reduced due to the disruption of postsynaptic elements of AMPA receptors (Bączyk et al., 2020a). While our data indicate that anodal polarisation increased the number of the available AMPA GluR4 subunits on the postsynaptic site of the Ia synapse, the significantly higher levels of available neurotransmitter—as indicated by an increased VGlut1 signal—might still oversaturate the receptors and not allow the full PP facilitation to develop. Still, one should note that the PP depression was not visible in a vast majority of the recorded MNs following anodal polarisation (Fig. 2C-C_1_) indicating that the recovery of postsynaptic receptors did indeed have a positive effect on MN physiology.

On the other hand, the impact of cathodal polarisation on the VGlut1 intensity fits well with the observed increase of the PP ratio in the Cathodal polarisation group. As the levels of the available neurotransmitter in the Ia synapse become reduced following cathodal polarisation (Fig. 4C, D), this leaves more postsynaptic receptors available for the second pulse of the test. Importantly, cathodal polarisation did not reduce the already low number of AMPA GluR4 subunits in the SOD1 animals. As a result, a marked PP facilitation was seen in the Cathodal polarisation group.

### Two weeks of tsDCS has a neglective impact on the activity of intracellular pathways and disease markers

Chronic activation of the PKA pathway in SOD1 animals leads to an increase in MN synaptic excitation and a decrease of misfolded SOD1 protein in spinal MNs (Bączyk et al., 2020a). As in our experiments, a significant increase in Ia EPSP amplitude was seen following two weeks of anodal tsDCS. We expected that this would provoke an increase in the activation of intracellular metabolic pathways and a reduction of misfolded SOD1 protein levels. However, this does not seem to be the case, as levels of pCREB and misfSOD1 were not affected by any type of polarisation. The lack of pCREB increase points to a fundamental disruption in the transmission of synaptic signals to intracellular pathways. Indeed, a “decoupling” of MN synaptic excitation from the intracellular pathways was recently reported by Grycz et al. (PS02-26PM-590 FENS Forum 2024). In their work, the authors applied vibration stimuli to the Achilles tendon and recorded the evoked compound EPSPs in the TS muscle, while simultaneously measuring the activity of activity-dependent intracellular pathways. It was found that the vibration-evoked EPSPs failed to increase pCREB levels in the SOD1 animals. Furthermore, when ampakine was used to increase the probability of opening the AMPA receptor in SOD1 MNs, no increase in the vibration-induced pCREB levels was found, despite a strong increase in the compound EPSP amplitude. This indicates that the activity of intracellular pathways is not directly connected to the MN synaptic excitation in SOD1 animals. Indeed, this result falls well in line with our investigation of tsDCS effects, where no increase in pCREB levels was seen despite a significant increase of maximal Ia EPSP amplitude. The lack of an increase in the pCREB levels is therefore a likely cause of the negligible effect of tsDCS on misfolded SOD1 levels.

### Study limitations

All care was taken to standardise the experimental conditions for each experiment. However, some factors potentially impacting the effects of the tsDCS application could not be avoided. The most important would be the inability to directly predict the spatial orientation of recorded neurons with respect to the applied electrical field. It has already been established in both in vitro experiments (Lafon et al., 2017; Rahman et al., 2013) and modelling studies (Elbasiouny and Mushahwar, 2007; Rattay, 1999) that radial and transverse electrical fields have different impacts on neuronal compartments—evoking their hyper- or depolarisation—depending on neuron spatial orientation. While spinal MN somas are located in the Rexed lamina IX of the ventral horns, the direct orientation of the axon initial segment and dendritic arborisation can vary between individual neurons (see Fig. 4). This means that some neurons might be preferentially affected by tsDCS.

Next, our investigations were confined to MNs innervating the TS motor pool. This approach was selected due to the substantial electrophysiological and molecular data profiles of these MNs already available from ours and collaborating laboratories. However, there is a significant difference between the rate of motor pool degeneration in ALS; pools innervating muscles containing predominantly fast muscle fibres (such as tibialis anterior, medial gastrocnemius) degenerate faster than those composed mainly of slow muscle fibres (e.g., soleus). It is possible that tsDCS has differential effects on flexor and extensor MNs, especially in the framework of different synaptic inputs to these motor pools.

Finally, tsDCS was applied to the lumbar segments of the spinal cord, therefore limiting the potential therapeutic effect of our intervention, because ALS affects all spinal motoneurons and motor cortical areas.

## Data availability statement

All data supporting the results of this study are included in the manuscript. The datasets generated during the current study are available from the corresponding author upon reasonable request. All raw recordings are stored in the private data repository at https://box.pionier.net.pl/ provided by the Poznan Supercomputing and Networking Center affiliated with the Institute of Bioorganic Chemistry of the Polish Academy of Sciences. Access to the raw data can be provided upon request.

## Acknowledgements

We thank Dr Stefano Antonucci for providing the Fiji macro to analyse the GluR4 cluster data, and Dr Simon Danner for his support in statistical analysis. We also thank Msc. Piotr Zawistowski and Msc. Bartosz Wasicki for participating in the long-term polarisation protocols, and Prof. Jakub Dalibur Rybaka and Msc. Tomasz Szczepański from WCZT for assistance in confocal microscopy of Ia synaptic contacts.

## Author contributions

All electrophysiological experiments and data analysis were performed at the Neurobiology Department of Poznan University of Physical Education. Imaging was done at the Wielkopolska Center for Advanced Technologies and Faculty of Materials Engineering and Technical Physics of Poznań Technical University imaging facilities. Conceptualization: M. Bączyk; methodology: M. Bączyk, K. Grycz; formal analysis: T. Jankowiak, M. Cholewiński, K. Grycz, E. Krok and M. Bączyk; investigation: T. Jankowiak, M. Cholewiński, K. Grycz, E. Krok, K. Kryściak and M. Bączyk; writing the original draft: T. Jankowiak, M. Cholewiński, K. Grycz, E. Krok, K. Kryściak and M. Bączyk; visualisation: K. Grycz, E. Krok and M. Bączyk; supervision: M. Bączyk; funding acquisition: M. Bączyk. All authors approved the final version of the manuscript. All authors agreed to be accountable for all aspects of the work in ensuring that questions about the accuracy or integrity of any part of the work are appropriately investigated and resolved. All authors qualify for authorship, and all those who qualify for authorship are listed.

## Supplementary Material

**Supplementary Figure 1.**
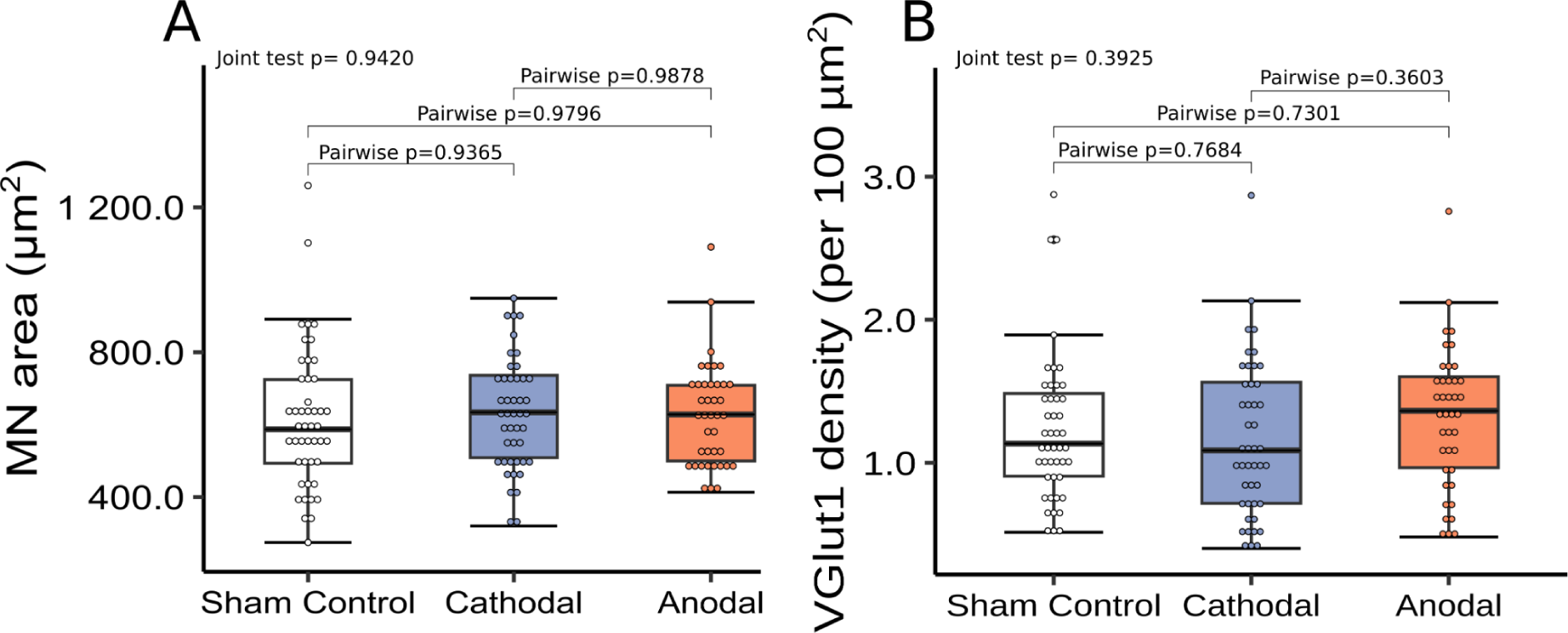
Effects of two-week polarisation on motoneuron size and synaptic coverage. Box plots showing data distribution of motoneuron area (**A**) and VgluT1 synaptic coverage (**B**). Box plots and statistical descriptions as in Figure 2. Note a lack of effect of polarisation on all analysed parameters.

**Supplementary Figure 2.**
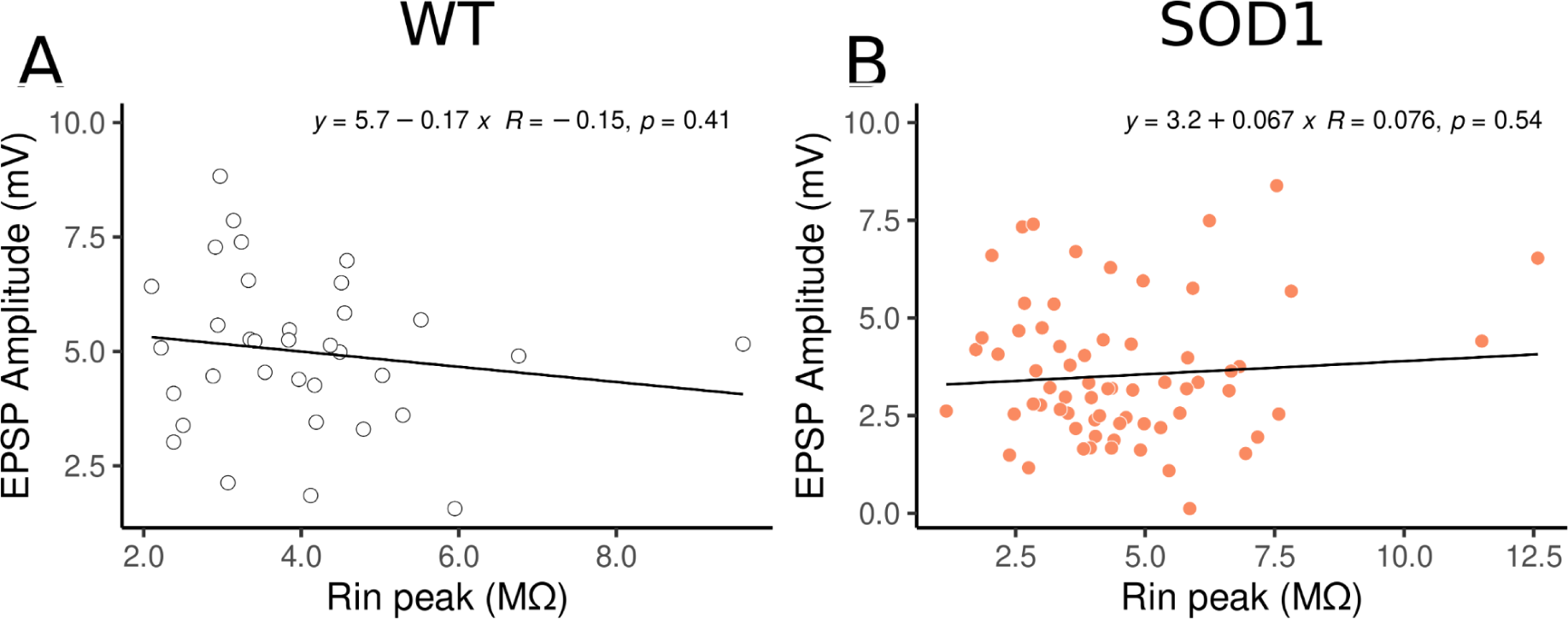
Correlation of maximal EPSP amplitude and peak input resistance in WT and SOD1 mice. Maximal Ia EPSPs recorded in triceps surae motoneurons (MNs) are plotted against cells’ peak input resistance for the WT (A) and SOD1 (B) mice. For each relationship, a linear trendline is plotted, and the Spearman correlation coefficient (R) with p-value is shown on the right of the trendline equation. On the plots, each data point represents a single MN. Plots use re-analysed data from Jankowiak et al. 2022.

**Supplementary Figure 3.**
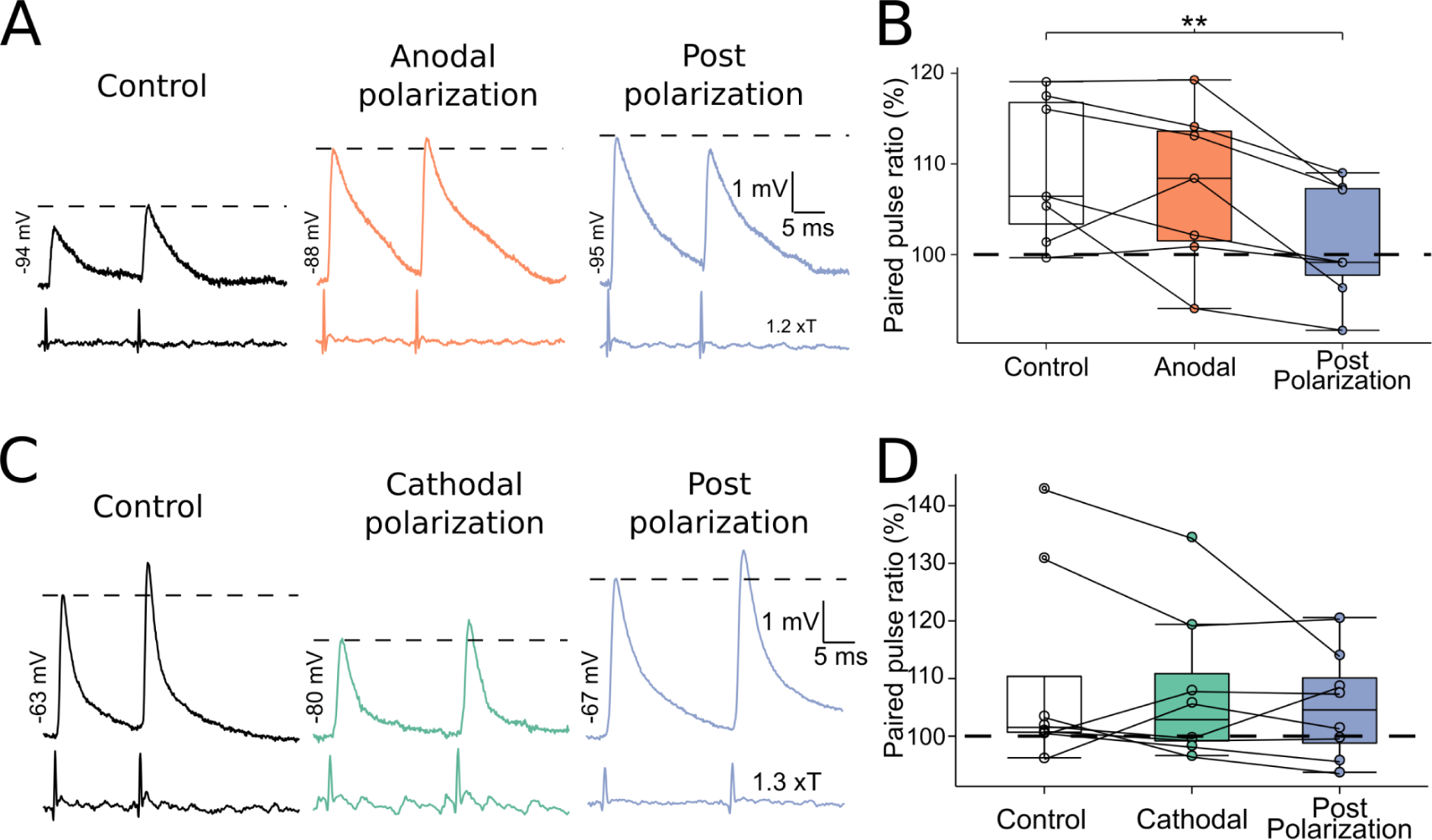
Acute tsDCS effects on the paired-pulse protocol in SOD1 mice. Intracellular recordings of paired-pulse response from the same triceps surae motoneuron (MN) evoked by Ia afferent stimulation, before, during and after anodal (**A**) or cathodal (**C**) polarisation. **B, D,** box plots showing data distribution of paired-pulse ratio following anodal and cathodal polarisation, respectively. Boxplot description as in Figure 2. The dashed lines show the amplitude of the first response from the paired-pulse pair (**A, C**) or the border between paired-pulse depression and paired-pulse facilitation (**B, D**). ** indicates a significant difference to Control at p<0.01. Each data point indicates a single MN from a single animal. Plots use re-analysed data from Jankowiak et al. 2022.

